# Chemical Stabilization of the HIV-1 Capsid Results in Efficient HIV-1 Reverse Transcription *in vitro*

**DOI:** 10.1101/2020.09.17.302315

**Authors:** Jordan Jennings, Jiong Shi, Janani Varadarajan, Parker J. Jamieson, Christopher Aiken

## Abstract

A defining activity of retroviruses is reverse transcription, the process during which the viral genomic RNA is converted into the double strand DNA required for virus replication. Reverse transcriptase (RT), the viral enzyme responsible for this process, was identified in 1970 by assaying permeabilized retrovirus particles for DNA synthesis *in vitro*. Such reactions are inefficient with only a small fraction of viral genomes being converted to full-length double strand DNA molecules, possibly owing to disruption of the structure of the viral core. Here we show that reverse transcription in purified HIV-1 cores is enhanced by the addition of the capsid-binding host cell metabolite inositol hexakisphosphate (IP6). IP6 potently enhanced full-length minus strand synthesis, as did hexacarboxybenzene (HCB) which also stabilizes the HIV-1 capsid. Both IP6 and HCB stabilized the association of the viral CA and RT proteins with HIV-1 cores. In contrast to the wild type, cores isolated from mutant HIV-1 particles containing intrinsically hyperstable capsids exhibited efficient reverse transcription in the absence of IP6, further indicating that the compound promotes reverse transcription by stabilizing the viral capsid. Our results show that stabilization of the HIV-1 capsid permits efficient reverse transcription in HIV-1 cores, providing a sensitive experimental system for analyzing the functions of viral and host cell molecules and the role of capsid disassembly (uncoating) in the process.

**IMPORTANCE:** HIV-1 infection requires reverse transcription of the viral genome. While much is known about the biochemistry of reverse transcription from simplified biochemical reactions, reverse transcription during infection takes place within a viral core. However, endogenous reverse transcription reactions using permeabilized virions or purified viral cores have been inefficient. Using viral cores purified from infectious HIV-1 particles, we show that efficient reverse transcription is achieved *in vitro* by addition of the capsid-stabilizing metabolite inositol hexakisphosphate. Enhancement of reverse transcription was linked to the capsid-stabilizing effect of the compound, consistent with the known requirement for an intact or semi-intact viral capsid for HIV-1 infection. Our results establish a biologically relevant system for dissecting the function of the viral capsid and its disassembly during reverse transcription. The system may also prove useful for mechanistic studies of emerging capsid-targeting antiviral drugs.

## INTRODUCTION

During retrovirus infection, the viral membrane fuses with the target cell, releasing the viral core into the cytoplasm. The core, consisting of a capsid shell surrounding the viral genome and its associated proteins, represents the functional viral payload. In the cell, the viral RNA genome is converted into a double strand DNA molecule by reverse transcription, producing the cis-acting viral sequences necessary for integration and subsequent gene expression. Reverse transcription is catalyzed by the viral reverse transcriptase enzyme (RT) and takes place in a ribonucleoprotein complex housed within the viral capsid. For HIV-1, pharmacological or genetic perturbations of the stability of the capsid typically result in impaired infectivity. Specifically, destabilization of the viral capsid leads to inefficient reverse transcription in target cells (1, 2) while hyperstabilization of the capsid inhibits nuclear entry and integration (3, 4). Similarly, premature capsid disruption in cells expressing restrictive TRIM5 proteins is associated with impaired reverse transcription (5, 6). Collectively, these studies have established that the integrity of the viral capsid is important for efficient HIV-1 reverse transcription in target cells.

HIV-1 reverse transcription occurs in a series of stages (reviewed in (7)). Synthesis of the minus strand is primed near the 5’ end of the genome by a tRNA, resulting in run-off synthesis of a short DNA molecule, the minus strand strong stop. Subsequently, this product anneals to the 3’ end of the genome and is extended, resulting in a ∼9 kb minus strand product. Plus-strand synthesis is primed by a small RNA remnant at the beginning of the U3 sequence, resulting in a short product that is then extended after annealing to the 3’ end of the minus strand. Subsequently, synthesis of the two viral long terminal repeat sequences (LTRs) is completed, resulting in a preintegration complex (PIC) that catalyzes integration of the nascent viral DNA into the target cell genome. A recent study suggests that nuclear entry precedes the completion of reverse transcription, suggesting that the core/reverse transcription complex responds to a specific nuclear signal or is activated during the process of nuclear entry (8). Nonetheless, it is known that active PICs containing two complete DNA ends can be recovered from the cytoplasm of acutely infected cells (9).

Inositol phosphates are abundant cellular metabolites that participate in a wide array of cell activities (reviewed in (10)). These highly charged small molecules include inositol (1,3,4,5,6) pentakisphosphate (IP5) and inositol hexakisphosphate (IP6). IP6 binds to numerous host cell proteins and regulates diverse biological processes, including chromatin remodeling (11), mRNA nuclear export (12, 13), platelet aggregation (14), prion propagation (15), and circadian rhythm (16). IP6 binds to the HIV-1 capsid *in vitro* and stabilizes the hexameric CA lattice. It associates with the center of the CA hexamer, forming ionic interactions with the six Arg18 side chains residing within the hexamer pore formed by the CA N-terminal domains (17, 18). In endogenous reverse transcription reactions with purified HIV-1 cores, the addition of IP6 protected the newly synthesized viral DNA from degradation by exogenously added DNaseI *in vitro*, suggesting that the viral capsid can provide an barrier to access of the viral genome (18). IP6 is incorporated into budding HIV-1 particles via an interaction with a distinct site in the assembling Gag lattice, near the CA-SP1 junction (17). It has been proposed that during maturation, IP6 is released upon proteolytic cleavage of Gag and subsequently associates with the mature capsid lattice and stabilizes it (17, 18). The critical importance of capsid stability in HIV-1 reverse transcription and infection, coupled with the relative biochemical instability of purified HIV-1 cores, makes this an appealing model. However, a role of IP6 in reverse transcription itself has not been established.

Although HIV-1 reverse transcription occurs efficiently in permissive target cells, *in vitro* assays of reverse transcription in permeabilized virions are typically inefficient, for unclear reasons. In these “endogenous reverse transcription” reactions, only a small fraction of viral genomes is converted into full-length double-strand DNA molecules. Such reactions have frequently relied on the addition of detergents or other membrane-disrupting agents to suspensions of concentrated virions, thus permitting access of dNTPs to the viral core (19, 20). The addition of detergents may compromise reverse transcription reactions by destabilizing the viral capsid, resulting in dissociation of RT from the template and its diffusion out of the viral core (21). HIV-1 reverse transcription complexes (RTCs) isolated from acutely infected cells generally lack substantial quantities of the CA protein (22), suggesting that the capsid dissociates during cell permeabilization. The apparent fragility of HIV-1 cores and reverse transcription complexes has hampered biochemical studies of early events in HIV-1 infection, specifically reverse transcription and the role of the viral capsid in this process.

To address this problem, we identified detergent-free experimental conditions in which HIV-1 cores purified from infectious virions undergo efficient reverse transcription. We show here that addition of the capsid-stabilizing cell metabolite IP6 markedly enhances the efficiency of reverse transcription by promoting the synthesis of full-length minus strand DNA. IP6 also stabilized the association of the CA and RT proteins with HIV-1 cores, suggesting that the effect was mediated by capsid stabilization. Our results are consistent with a model in which the viral capsid promotes retention of a sufficient concentration of RT in association with the viral genome to ensure completion of reverse transcription.

## RESULTS

### Establishment of the endogenous reverse transcription (ERT) reaction using purified HIV-1 cores

In an effort to improve the efficiency of ERT, we incubated samples of purified HIV-1 cores with dNTPs *in vitro* and analyzed the products by quantitative PCR. For this purpose, we purified HIV-1 cores by a method involving ultracentrifugation of concentrated virions through a layer of Triton X-100 detergent into a sucrose density gradient. Under these conditions, the virions are exposed to the detergent for only a brief time, thus preserving the integrity of the viral core. During centrifugation, HIV-1 cores sediment into the gradient, resulting in removal of the detergent. Cores were detected in gradient fractions by p24 ELISA for the CA protein (Fig. 1A), by assay for RT activity (Fig. 1B), and by negative-stain electron microscopy (Fig. 1C).

**FIG 1.**
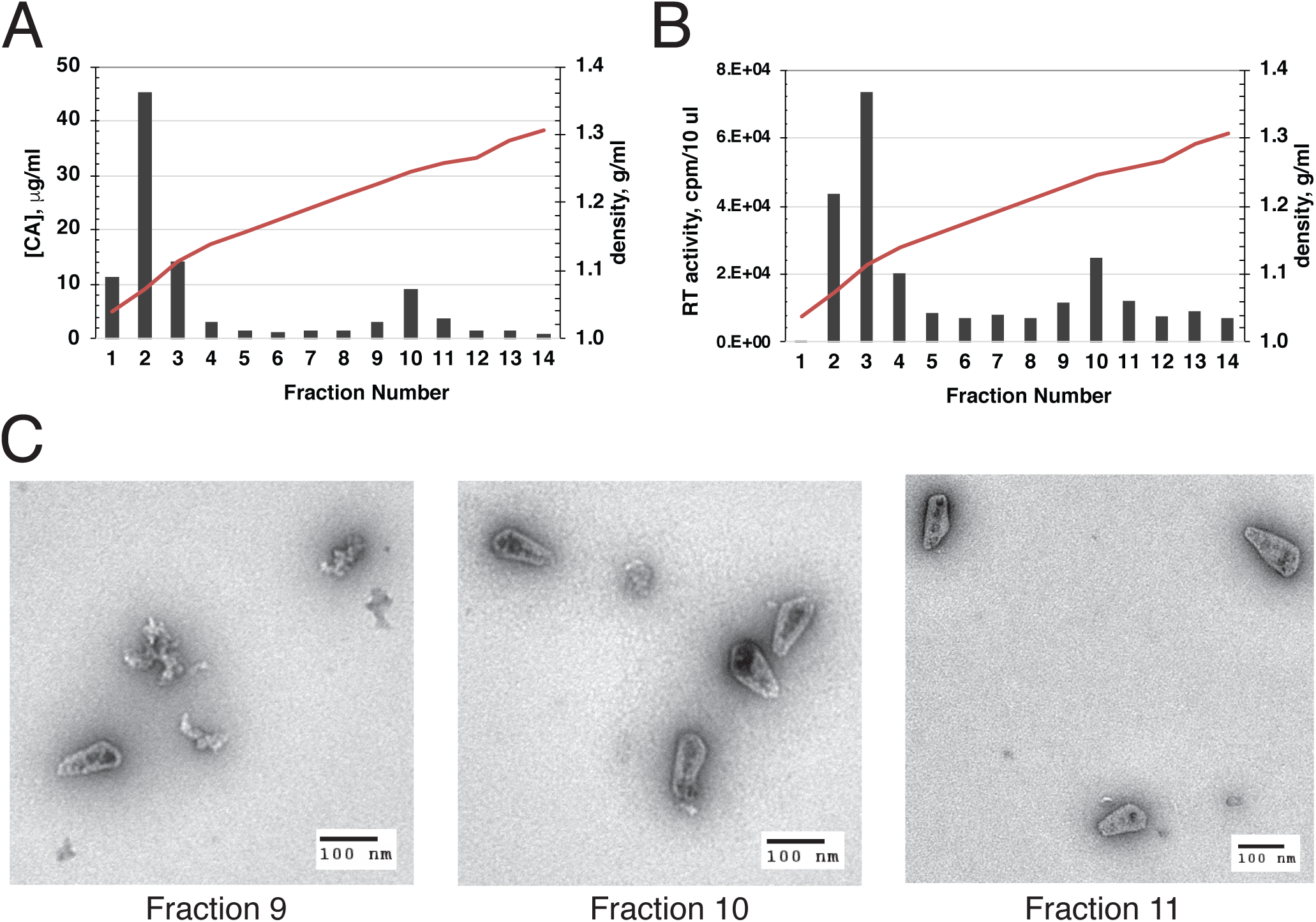
Characterization of purified HIV-1 cores. HIV-1 cores were isolated from concentrated virions by sucrose density gradient sedimentation. Gradient fractions were assayed for (A) CA protein and (B) RT activity. Panel C shows electron micrographs from negative stained samples of the three gradient fractions containing HIV-1 cores.

In initial studies, we attempted to improve the reaction efficiency by varying the temperature, reaction time, and pH. We also tested polyethylene glycols which have been previously reported to stimulate reverse transcription *in vitro* (23), and tested the effects of adding bovine serum albumin (BSA). After extended incubation at 37°C, the DNA products were purified and quantified for sequences corresponding to various stages of reverse transcription. An example of the results obtained in this type of experiment is shown in Fig. 2. Quantitative PCR (qPCR) analysis demonstrated that some of these parameters resulted in modest increase in overall ERT efficiency (based on the ratio of 2_nd_ strand transfer products to minus strand strong stop molecules). In particular, the addition of bovine serum albumin appeared beneficial. Nonetheless, based on the quantitative analysis of each stage of reverse transcription, the overall efficiency of the reactions appeared to be limited by a marked attenuation in full-length minus strand synthesis. By contrast, both strand transfer events were relatively efficient. These results indicated that the cores initiated reverse transcription but were unable to synthesize the complete minus strand. Of note, the reactions required prolonged incubation times, suggesting the possibility that the viral capsid dissociated during the incubation, resulting in loss of RT, as previously observed (1).

**FIG 2.**
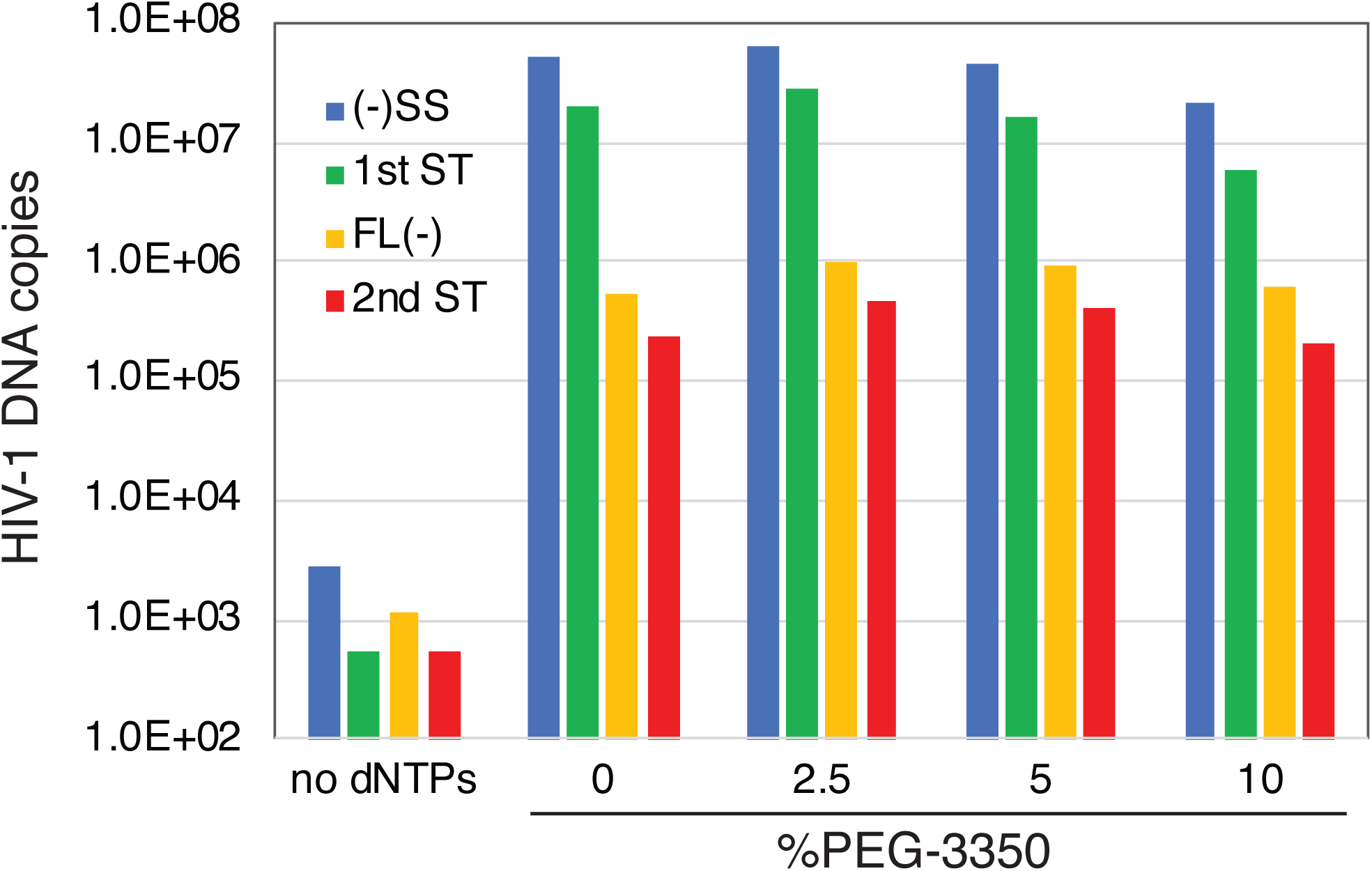
Representative results from an ERT experiment. HIV-1 cores were incubated for 16h in a preliminary ERT reaction buffer containing the indicated concentrations of polyethylene glycol 3350. DNA was purified in the reactions and analyzed by qPCR for the indicated products of reverse transcription. This experiment was one of many early attempts to improve the efficiency of ERT.

### IP6 promotes efficient minus strand synthesis in ERT reactions

The cell metabolite IP6 was recently reported to bind to the HIV-1 capsid *in vitro* and to promote the assembly of recombinant CA protein into capsid-like structures (17, 18). Adding IP6 to permeabilized HIV-1 particles resulted in protection of newly synthesized viral DNA from degradation by DNaseI. By stabilizing the viral capsid, IP6 may prevent access of the nuclease to the nascent viral DNA. However, in that study, addition of IP6 did not enhance ERT when performed in the absence of DNaseI. To test whether capsid stabilization by IP6 can enable efficient ERT, we performed reactions in the presence of a range of IP6 concentrations (Fig. 3). We observed a marked enhancement of full-length minus strand synthesis and increased overall efficiency of the reaction in the presence of low micromolar concentrations of IP6. These results indicate that IP6 increases the efficiency of ERT.

**FIG 3.**
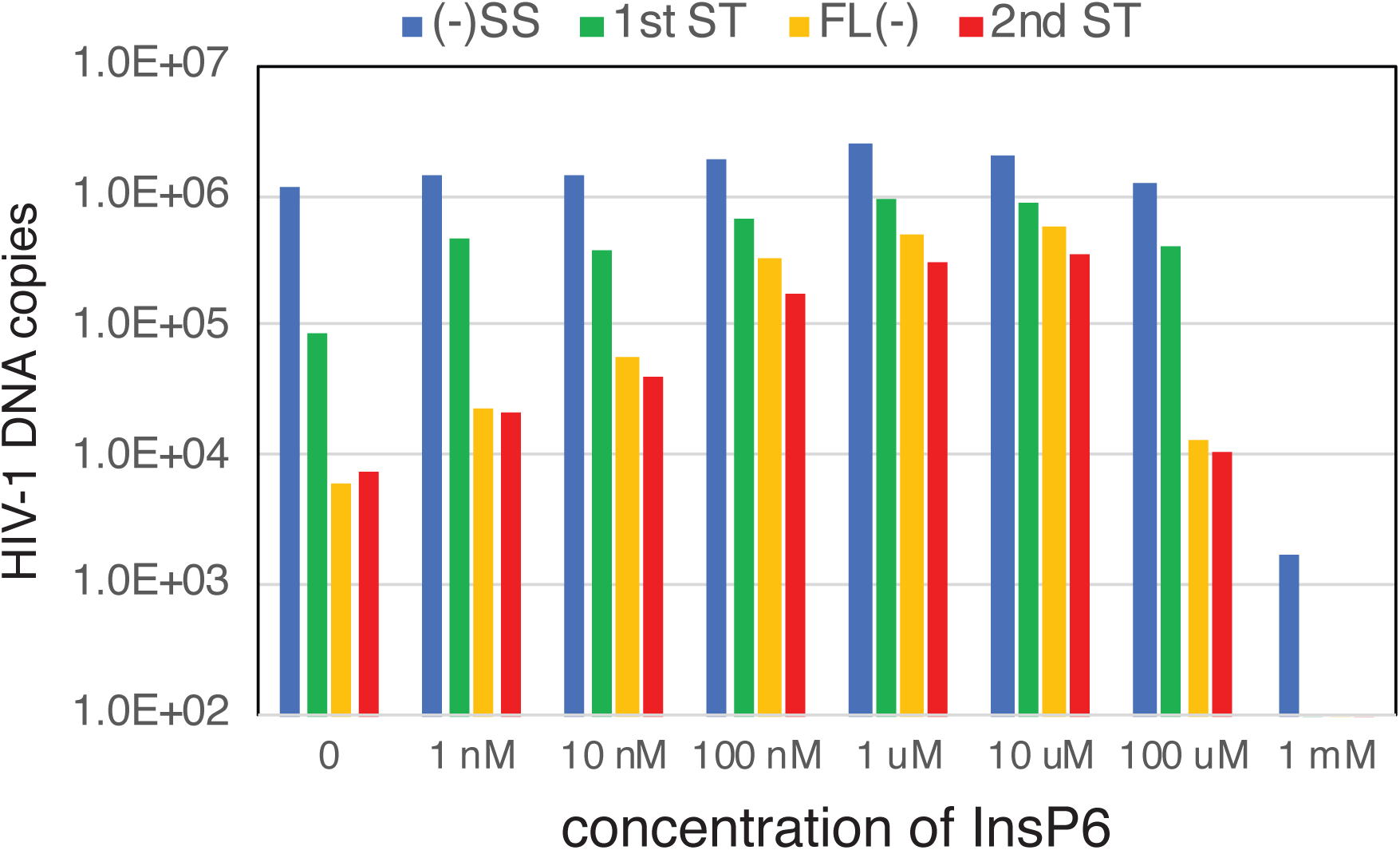
IP6 markedly stimulates ERT in vitro by enhancing minus strand synthesis. ERT reactions containing the indicated concentrations of IP6 were incubated at 37°C for 16h, and DNA products were purified and quantified.

Following this observation, we optimized several parameters in reactions containing 10 μM IP6, including NaCl and MgCl_2_ concentrations and pH. We thus identified conditions for efficient ERT: 10 mM Tris-HCl pH 7.6, 150 mM NaCl, 2 mM MgCl_2_, 1 mg/ml BSA, 0.5 mM DTT, and 10 μM IP6, with incubation at 37°C for 16h. Under these conditions, we reproducibly observed conversion of 35 to 40% of the minus strand strong stop products into molecules that also contained HIV-1 sequences that are synthesized only after the second-strand transfer step (Fig. 4). This was the maximum ERT efficiency we observed in multiple experiments with different preparations of cores.

**FIG 4.**
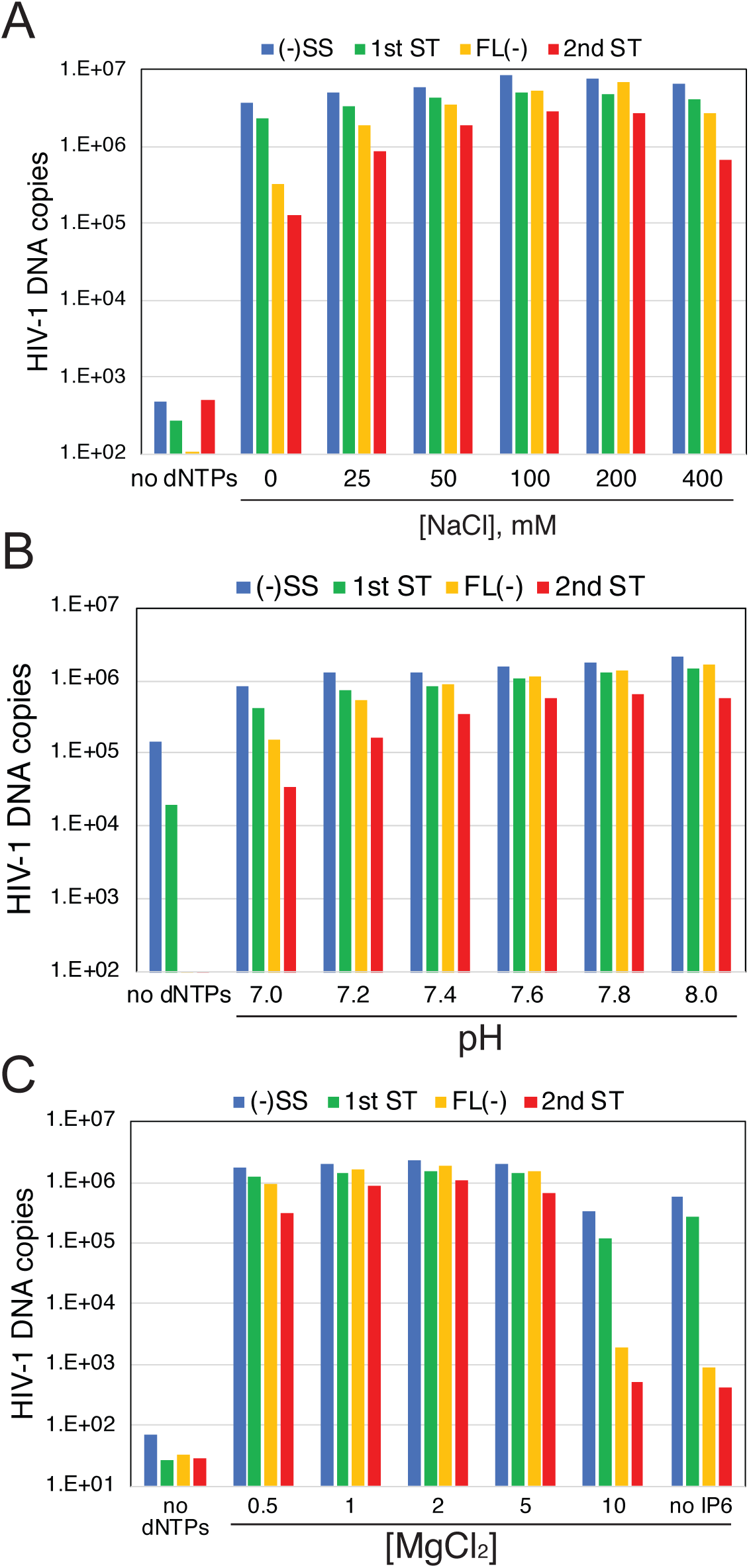
Optimization of the ERT reaction. ERT reactions containing 10 μM IP6 and the indicated conditions were incubated and analyzed for HIV-1 DNA products. The variables were: (A) NaCl concentration, (B) pH, in reactions containing 150 mM, and (C) MgCl_2_ concentration, in reactions at pH 7.6 with 150 mM NaCl.

### Kinetics of the ERT reaction

We also analyzed the rates of product formation in ERT reactions under the optimized conditions. Minus strand strong stop and 1_st_ strand-transfer products were synthesized rapidly in reactions containing IP6, reaching half-maximal values within 2 h (Fig. 5A). By contrast, synthesis of full-length minus strand and 2_nd_ strand transfer products required 4 to 8 h to reach half-maximal values. In reactions lacking IP6, both early product species were also 50% complete after 2 h, but declined slightly, suggesting the possibility of partial degradation (Fig. 5B). Late products were produced at 2 h in reactions lacking IP6 and declined thereafter. Late products were then slightly increased at the 16h time point, suggesting that degradation may compete with ongoing synthesis. While IP6 stimulated the synthesis of early products up to ten-fold, the effect on late stage reverse transcripts was profound, enhancing product accumulation by several thousand-fold (Fig. 4, compare panels A and B). PCR quantification of the 16h ERT products generated in the absence of IP6 using primers spanning the genome revealed that extension of the minus strand was impaired with few DNA products longer than 3 kb accumulating in the reactions (Fig. 5C).

**FIG 5.**
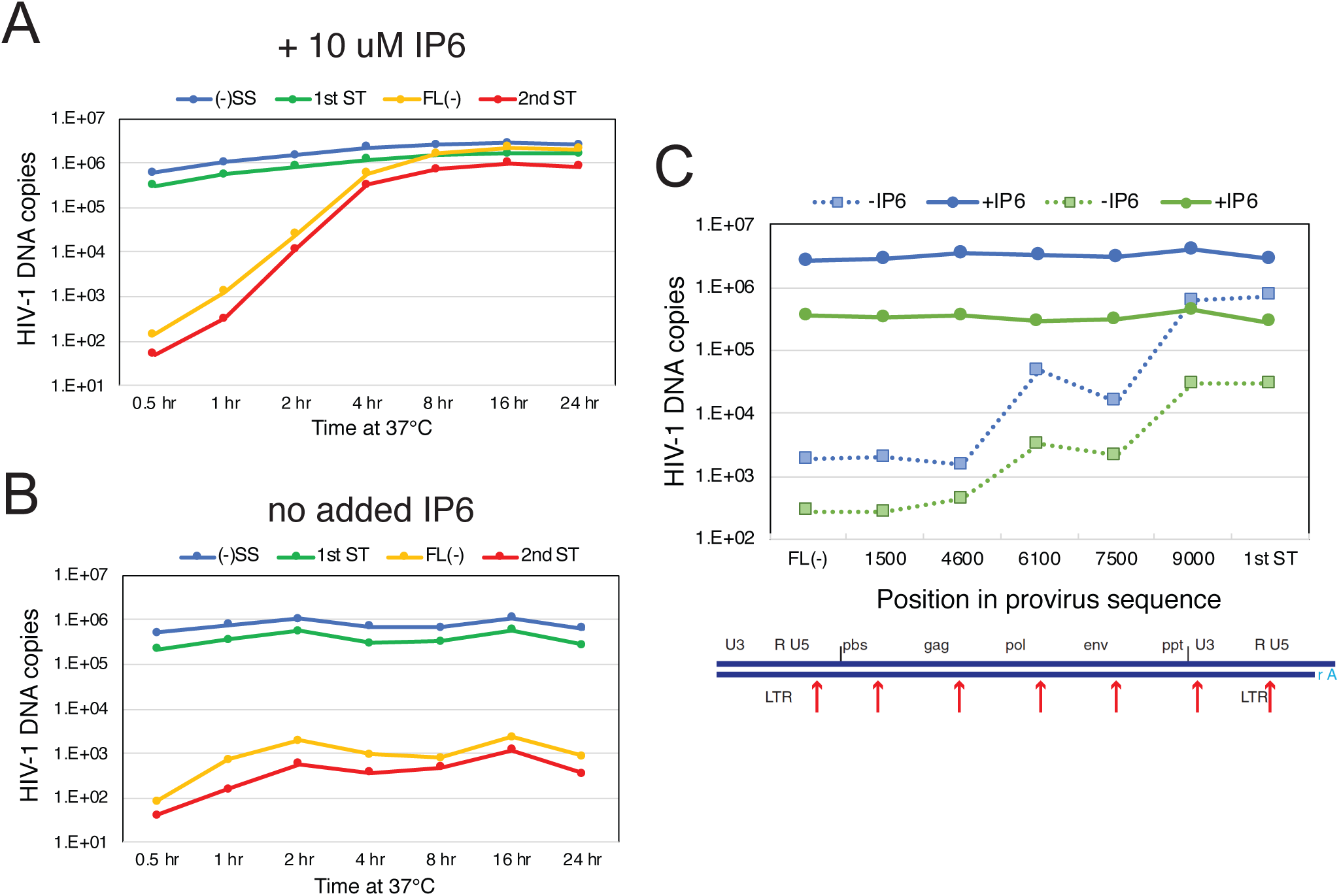
Time course of ERT in the presence and absence of IP6. ERT reactions containing and lacking 10 μM IP6 were incubated for the indicated time periods and subsequently analyzed for various HIV-1 DNA products by qPCR. (A) reactions containing IP6; (B) reactions lacking IP6. Values represent averages of duplicate ERT reactions. (C) Quantification of products in 16h reactions for sequences spanning the viral genome in reactions containing and lacking IP6. The blue and green symbols represent values from pairs of ERT reactions from two different experiments. Dashed lines connect values from ERT reactions lacking IP6. Results shown in this figure are from one of two independent experiments.

### Hexacarboxybenzene stimulates ERT

We also tested the synthetic hexavalent compound hexacarboxybenzene (HCB) in ERT reactions, owing to a previous report that HCB stabilizes HIV-1 cores *in vitro* (24). Addition of HCB markedly enhanced minus strand synthesis at an optimal concentration of ∼100 μM, resulting in efficient ERT (Fig. 6A). However, HCB inhibited all stages of ERT when present at a concentration of 1 mM (Fig. 6B), consistent with a previous study reporting inhibition of ERT inhibition by 20 mM HCB (24).

**FIG 6.**
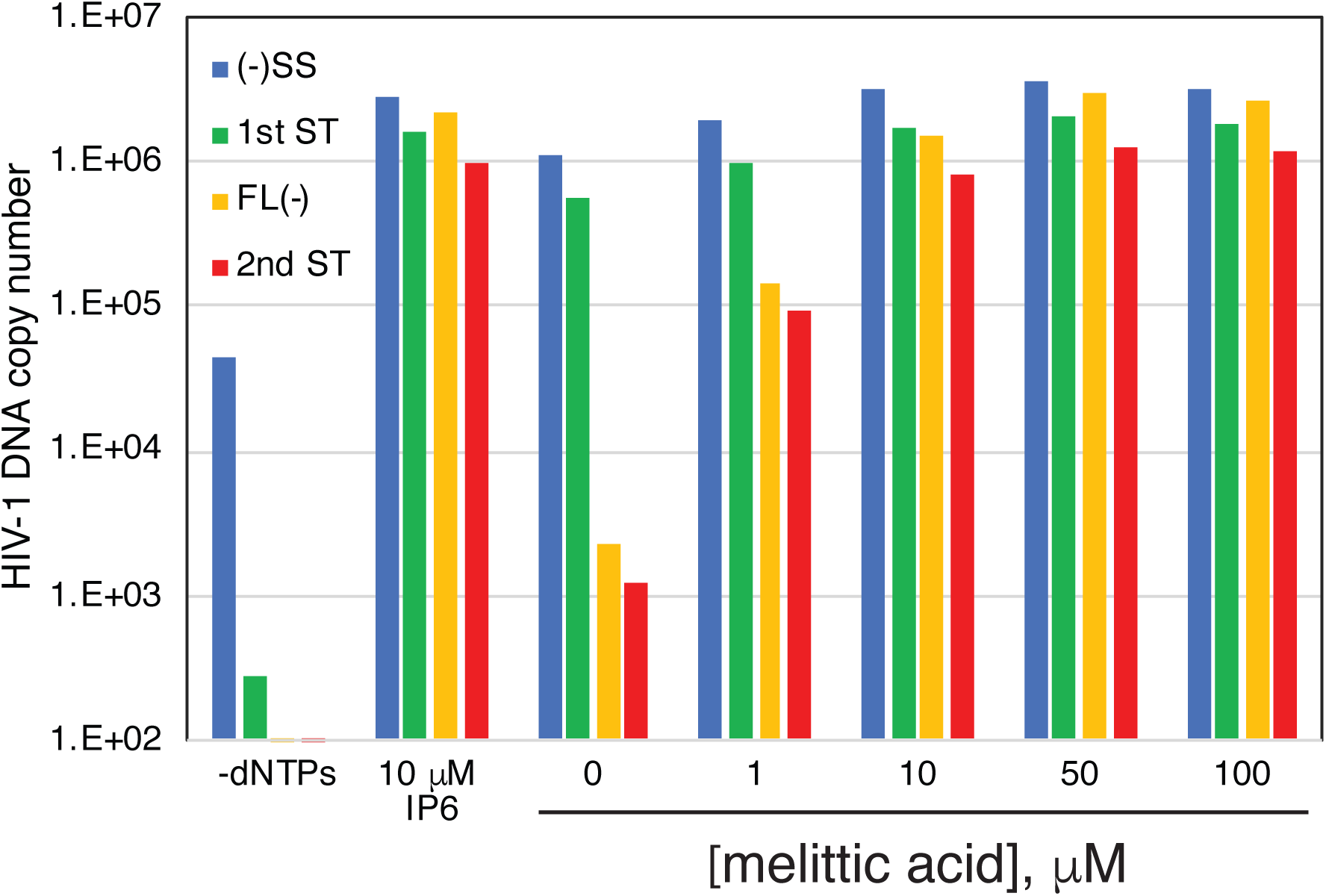
The capsid-stabilizing compound HCB also stimulates ERT. Reactions were containing the indicated concentrations of HCB were incubated for 16h at 37°C, and the products were analyzed by qPCR. Results shown are representative of two independent experiments.

To determine whether IP6 and HCB stabilize the HIV-1 capsid during ERT, we analyzed the quantity of HIV-1 CA and RT released from viral cores in reactions containing these compounds. Following a 6 h reaction time, the reactions were diluted with cold buffer and subjected to ultracentrifugation to separate the soluble proteins from those which remained core associated. Quantification of the fraction of the total CA present in the pellets revealed higher levels of pelletable CA protein in reactions containing IP6 or HCB, indicating that the viral capsid was stabilized by the compound (Fig. 7). Similarly, assays of RT activity in the supernatants and pellets showed that IP6 and HCB increased the levels of pelletable RT in the reactions. Collectively, these results suggest that the enhancing effects of IP6 and HCB on ERT result from stabilization of the viral capsid.

**FIG 7.**
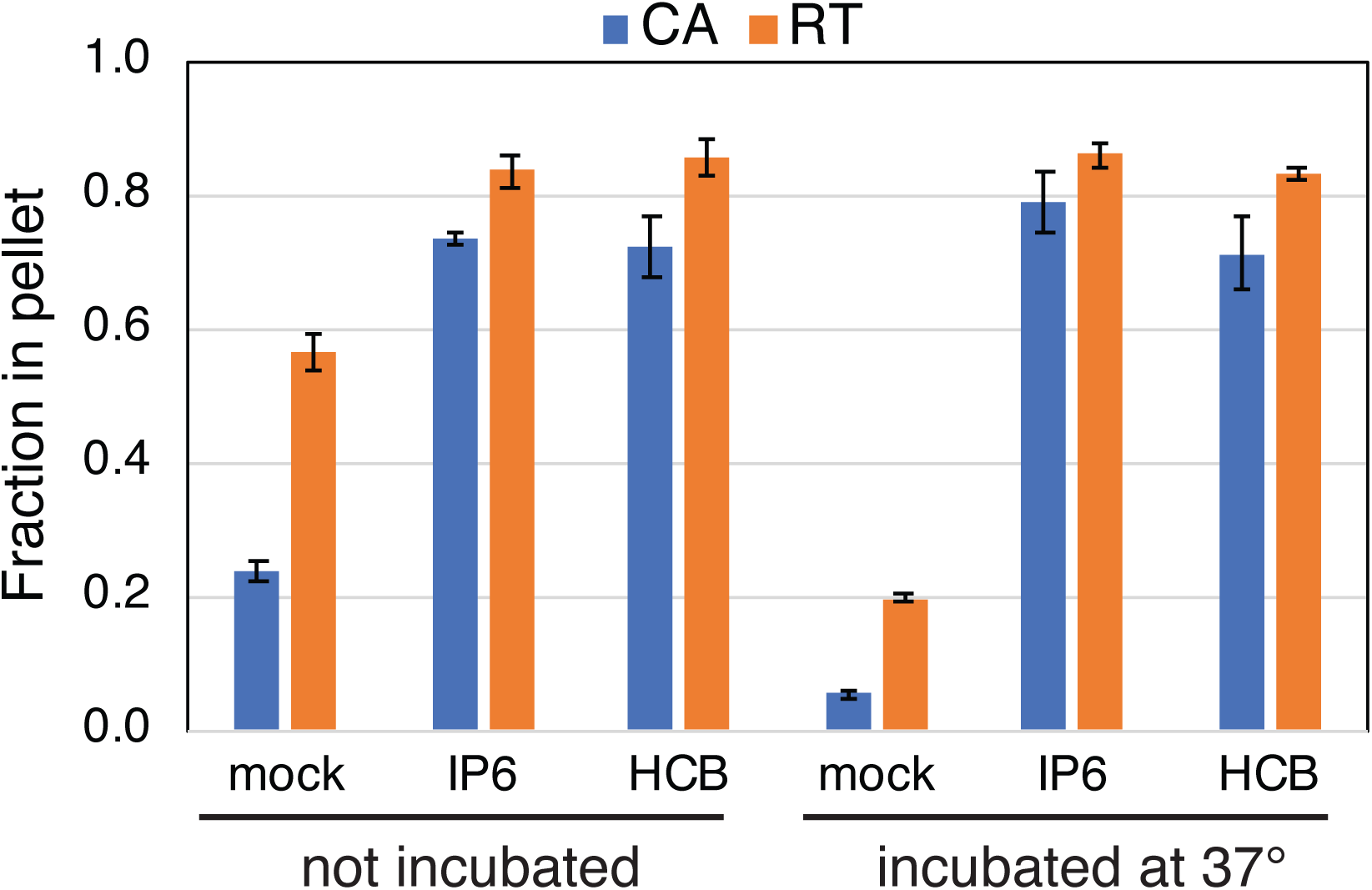
IP6 and HCB stabilize viral cores during reverse transcription. ERT reactions containing no additive, 10 mM IP6, or 100 mM HCB not incubated or incubated at 37°C for six hours. Reactions were diluted tenfold with reaction buffer, and the cores were pelleted by ultracentrifugation and the pellets and supernatants analyzed for CA and RT activity. Shown is the fraction of the total CA and RT activity in the pellets. The values shown are the average values of duplicate reactions from one of two independent experiments, which showed similar outcomes.

### Cores from an HIV-1 mutant with a hyperstable capsid undergo efficient ERT in the absence of added capsid stabilizers

We also asked whether genetic stabilization of the HIV-1 capsid affects the dependence of ERT on IP6. For this purpose, we isolated cores from HIV-1 particles containing the capsid-stabilizing CA substitution E45A. This mutant is competent for reverse transcription in target cells but is poorly infectious, owing to impaired nuclear entry and integration (4). Cores from the mutant are hyperstable *in vitro*, as inferred from their increased levels of core-associated CA and slower dissociation of CA during incubation at 37°C (1). In reactions containing IP6, the mutant cores exhibited efficient ERT, as did those from the wild type (Fig. 8). In reactions lacking IP6, E45A cores produced approximately 20% of the late stage products relative to parallel reactions containing IP6. This is in stark contrast to reactions with wild type cores, in which synthesis of late products was less than 0.1% of that observed in reactions containing IP6. These results indicate that E45A mutant cores are capable of synthesizing substantial quantities of late reverse transcripts in the absence of capsid-stabilizing agents, further indicating that the ERT-stimulating activity of IP6 results from stabilization of the viral capsid.

**FIG 8.**
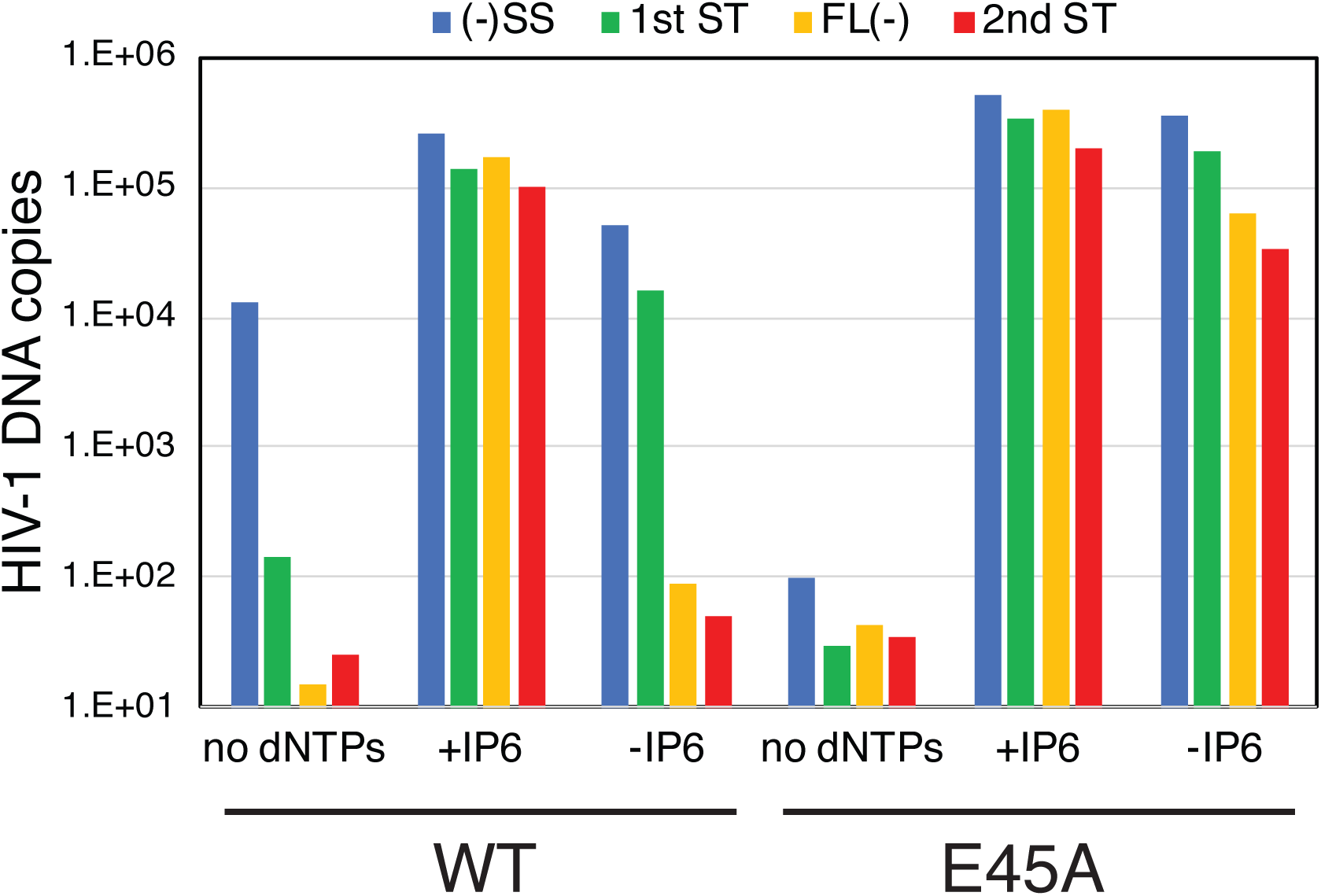
Cores from the E45A HIV-1 mutant, which contains a hyperstable capsid, are less dependent on IP6 or HCB for ERT. Cores were purified from HIV-1 particles that had been produced by transfection of 293Tcells. ERT reactions were performed with and without added IP6 or HCB.

### The capsid-targeting compound PF74 inhibits ERT in a concentration-dependent manner

Finally, we tested the effects of the capsid-targeting antiviral compound PF74. PF74 binds to a site in the CA hexamer that is distinct from that bound by IP6 (25, 26). When present during HIV-1 infection at concentrations of 10 μM and above, PF74 inhibits reverse transcription and destabilizes the viral capsid (27). Addition of 10 μM PF74 inhibited ERT in reactions containing IP6 (Fig. 9), further linking capsid function to ERT efficiency. By contrast, addition of 1 μM PF74 did not substantially inhibit ERT, consistent with previous reports that at low concentrations PF74 inhibits HIV-1 infection by affecting nuclear entry and integration (28-30). These results further support a role of the viral capsid in ERT.

**FIG 9.**
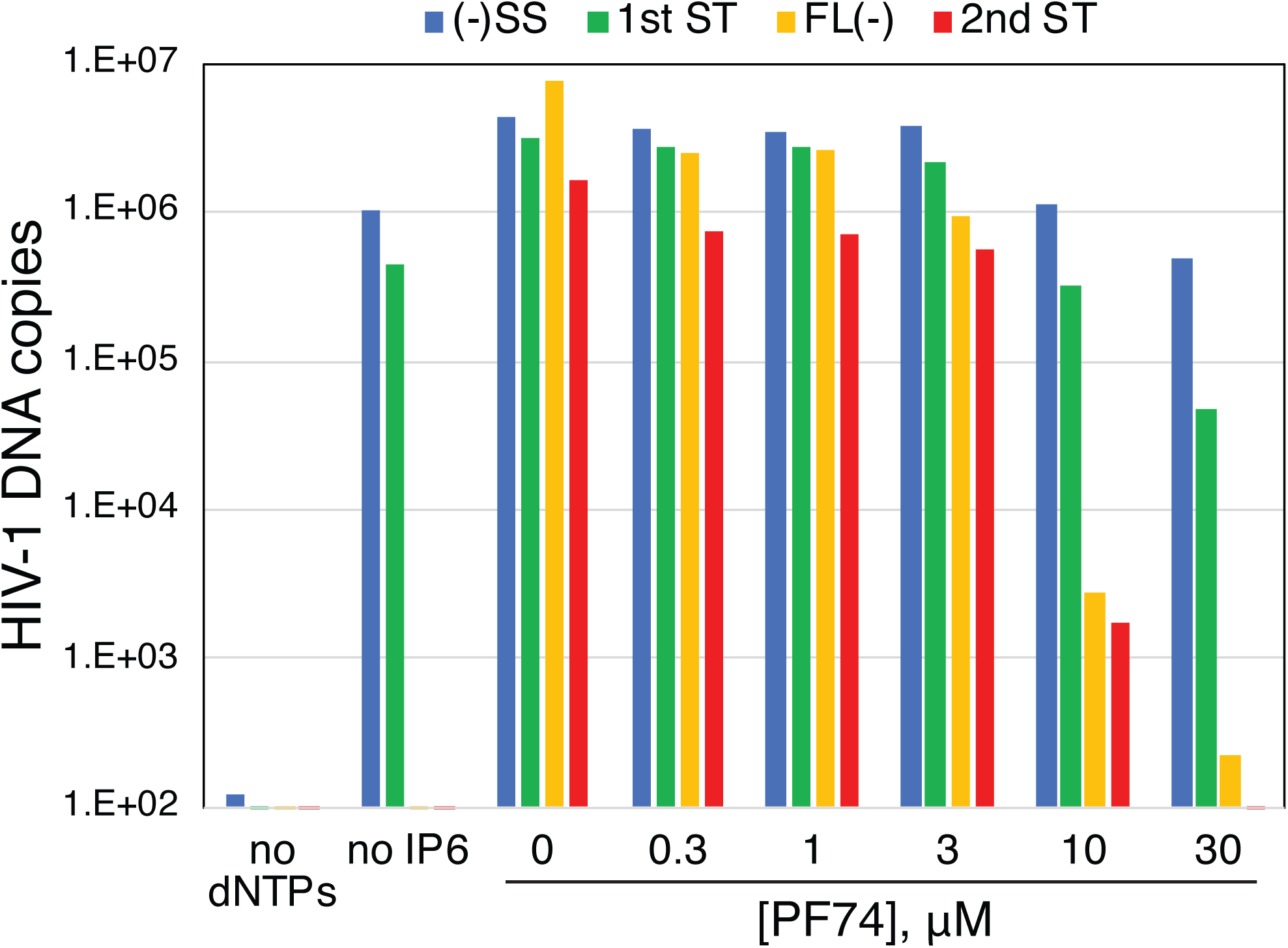
The capsid-targeting antiviral compound PF74 inhibits ERT. Optimized ERT reactions were containing indicated concentrations of PF74 were incubated for 16h. DNA products were purified and analyzed for HIV-1 sequences by qPCR. These results are representative of 3 independent experiments.

## DISCUSSION

In this study, we observed that addition of IP6 stabilizes HIV-1 cores and markedly enhances the efficiency of reverse transcription *in vitro*. Mallery and coworkers had previously shown that IP6 stabilizes purified HIV-1 cores and protects the products of ERT from degradation by added DNaseI, suggesting that the capsid provides a barrier to access to the synthesized DNA (18). In that study, IP6 did not appear to substantially alter the quantity of DNA products in the absence of added DNase. We also observed no enhancing effect of 1 mM IP6 on ERT. By contrast, addition of low concentrations of IP6 resulted in a nearly quantitative conversion of the initial minus strand products to full-length molecules. The observed enhancement resulted from a thousand-fold increase in minus strand synthesis together with modest enhancements at both strand transfer steps. We conclude that IP6 mainly promotes the completion of minus strand synthesis. The maximum efficiency of these reactions was ∼40% based on the ratio of 2_nd_ strand transfer products relative to minus strand strong stop molecules. Our observations suggest that purified cores can, under appropriate conditions, undergo efficient reverse transcription *in vitro* in the absence of added host proteins.

IP6 and the synthetic compound HCB also stabilized the association of CA and RT with viral cores, as previously observed in imaging studies of permeabilized virions. Thus, a plausible mechanistic conclusion is that the observed enhancement of ERT resulted from capsid stabilization, because most HIV-1 mutants containing intrinsically unstable capsids are impaired for reverse transcription in target cells (1). We also observed that mutant HIV-1 cores with hyperstable capsids synthesized substantial quantities of full-length minus strand DNA in reactions lacking IP6. Finally, we observed that the capsid-destabilizing HIV-1 inhibitor PF74 inhibited ERT even in the presence of IP6. Together, these observations support a capsid-stabilization mechanism for enhancement of ERT by IP6.

How might stabilization of the viral capsid help promote the completion of reverse transcription? In our experiments, the addition of IP6 resulted in a profound enhancement of minus strand DNA synthesis with lesser effects on the other steps that were quantified, including both strand transfers. In earlier work, our group showed that spontaneous uncoating of purified HIV-1 cores *in vitro* is characterized by dissociation of both CA and RT from the viral core (1). We suggest that the capsid acts as a container to retain RT during synthesis of the long (∼9 kb) minus strand DNA. HIV-1 particles are estimated to contain approximately 50 molecules of RT (7), with about 20% of the enzyme copurifying with cores (Fig. 1B). While completion of reverse transcription is theoretically possible with a single molecule of the enzyme, the relatively low processivity of the enzyme together with its frequent pausing suggest that RT must repeatedly rebind the template in order to synthesize full-length viral DNA. By preserving the association of RT with the core, the viral capsid may ensure that RT is maintained at a sufficient concentration to allow completion of the reaction. This “container model” does not exclude additional possible functions of the capsid in reverse transcription, such as providing a scaffold on which the reaction occurs.

IP6 is a natural metabolite that is present in mammalian cells at concentrations ranging from 40 to 90 μM (31), coinciding well with the ERT-enhancing effects we observed in the present study. Therefore, one could expect that HIV-1 infection would be strongly depending on target cell levels of IP6 and/or the related metabolite IP5. However, a recent study reported that ablation of cell expression of host proteins that synthesize these compounds does not appreciably influence their susceptibility to HIV-1 infection (32). These studies were performed in a transformed epithelial cell line, and it will be important to determine whether these inositol phosphates influence HIV-1 infection of physiologically relevant target cells. Additionally, purified HIV-1 cores appear to contain substantial quantities of bound IP6 (18), suggesting that the viral capsid may be stabilized by the IP6 it captures during assembly.

Reverse transcription is thought to promote HIV-1 uncoating in target cells, and it leads to physical changes in HIV-1 cores *in vitro* (33-35). The ERT reaction described herein mimics the kinetics of reverse transcription observed during synchronous HIV-1 infection of target cells (36), albeit without the initial lag phase, which presumably results from the requirement for virus fusion. Moreover, the reaction appears to be highly synchronous, with early DNA products appearing rapidly and the subsequent products accumulating only after a substantial delay. The experimental system described herein should enable structural and biochemical studies of native HIV-1 reverse transcription complexes and analysis of the role of the viral capsid in the process, including high-resolution analysis of the effects of reverse transcription on the structure of the viral core. ERT reactions with purified cores may also permit the generation of substantial quantities of pure and active HIV-1 PICs *in vitro* for biochemical studies of HIV-1 integration.

## Materials and Methods

### Chemicals, cells, and plasmids

Inositol hexakisphosphate was purchased as a 1.1M solution from TCI America (cat. No. P0409). D-myo-Inositol-1,3,4,5,6-pentakisphosphate (ammonium salt) was purchased from Cayman Chemical (cat. No. 10009851). Tris, mellitic acid, and polyethylene glycol 3350 were purchased from Sigma. Deoxynucleoside triphosphates (dNTPs) were purchased from New England Biolabs as 100 mM solutions. Bovine serum albumin was purchased from RPI (cat. No. A30075). The following antibodies were used for probing immunoblots: 183-H12-5C (NIH AIDS Research and Reference Program (37), used at 4 μg/ml); HIV-1 NC (polyclonal goat serum, from Dr. Robert Gorelick, used at a 1:1000 dilution); HIV-1 RT; HIV-1 IN (polyclonal rabbit serum, received from Dr. Alan Engelman, used at a 1:5000 dilution). The IR dye-conjugated polyclonal secondary antibodies were purchased from Li-Cor, Inc. The antiviral compound PF74 was synthesized and purified by the Chemical Synthesis Core of the Vanderbilt University Institute for Chemical Biology. For qPCR, the Maxima SYBR Green/ROX qPCR Master Mix (2X) from ThermoScientific (Cat. #K0233) was used. Custom oligodeoxyribonucleotides were purchased from Integrated DNA Technologies.

### Purification and analysis of HIV-1 cores

For most experiments, HIV-1 cores were purified from 200 ml of virus particles collected from infected MT4 cells. Cultured MT4 cells (1 x 10_7_) were pelleted and resuspended in 50 ml of medium. Cultures were inoculated wild type HIV-1 particles produced by transfection of 293T cells with the wild type R9 proviral construct. A quantity of the virus stock, corresponding to approximately 5 μg of p24, was added to the MT4 cultures with DEAE-dextran at a final concentration of 20 μg/ml. The following day, the cells were pelleted and resuspended in 200 ml of fresh culture medium. The cultures were examined daily signs of virus-induced cytopathicity, and at day 4-to 5 after inoculation, the cultures were centrifuged to remove cells and cell debris. The virus-containing culture supernatants were clarified by filtration and concentrated by ultracentrifugation at 32,000 rpm in a Beckman SW32.1Ti rotor at 4°C for 3h. Concentrated virions were resuspended in a total volume of 0.5 ml.

For experiments shown in Figure 8, HIV-1 cores were purified from virions produced by transfection of 293T cells, as previously described (38), with the following modifications. Four million 293T cells were transfected with 10 μg of R9 and R9.E45A plasmid DNAs using polyethyleneimine (39). The next day, cultures were washed and replenished with fresh medium. The following day, the culture supernatants were collected, clarified by filtration through a 0.45 μm vacuum filtration device, treated with 20 μg/ml DNaseI and 10 mM MgCl_2_ at 37°C for one hour to eliminate residual carryover plasmid DNA, then concentrated for purification of viral cores.

To purify cores, concentrated HIV-1 particles were subjected to centrifugation through a layer of Triton X-100 (1% vol/vol) into a linear sucrose density gradient, as previously described (38). Fractions (1 ml) were collected and assayed for CA protein by p24 ELISA and RT activity. The dense fractions corresponding to HIV-1 cores typically contained approximately 15% of the total CA protein in the gradients. Fractions containing HIV-1 cores were aliquoted, flash frozen in liquid nitrogen, and stored at -80°C for ERT reactions and other assays.

Negative stain electron microscopy of purified cores was performed by applying 3-5 μl samples of core preparations directly to glow-discharged carbon-coated copper EM grids. Following one minute of adherence, the excess liquid was removed by wicking, and the grids were inverted for one minute onto two consecutive droplets containing uranyl formate stain, followed by two consecutive inversions onto water droplets. The liquid was carefully removed by wicking, and the grids were air-dried and imaged in a Morgagni electron microscope. Images were captured using a CCD camera.

For immunoblot analysis of HIV-1 cores, 15 μl volumes of the gradient fractions were separated by electrophoresis on precast 4-20% polyacrylamide gels (Genscript). The proteins were transferred to nitrocellulose membrane, blocked with a solution of nonfat dry milk in TBST, and sequentially probed with antiserum to HIV-1 RT, IN, and CA proteins. Following probing of the blots with the appropriate IR dye-conjugated secondary antiserum, the corresponding HIV-1 protein bands were visualized by scanning the blots in a Li-Cor Odyssey imager.

### Assays of ERT using purified HIV-1 cores

ERT reactions were performed in 50 ul volumes containing 10 mM Tris-HCl pH 7.6, 150 mM NaCl, 2 mM MgCl_2_, 0.5 mM dithiothreitol, 0.1 mM each of 4 dNTPs, 1 mg/ml bovine serum albumin, and various concentrations of additives including IP6, mellitic acid, PEG-3350, and PF74. Reactions were normally incubated at 37°C for 16h, after which the DNA was extracted using a silica gel-based method (40) and eluted in water. The products were quantified by qPCR with SYBR green detection using the primers listed in Table 1:

**Table 1.**
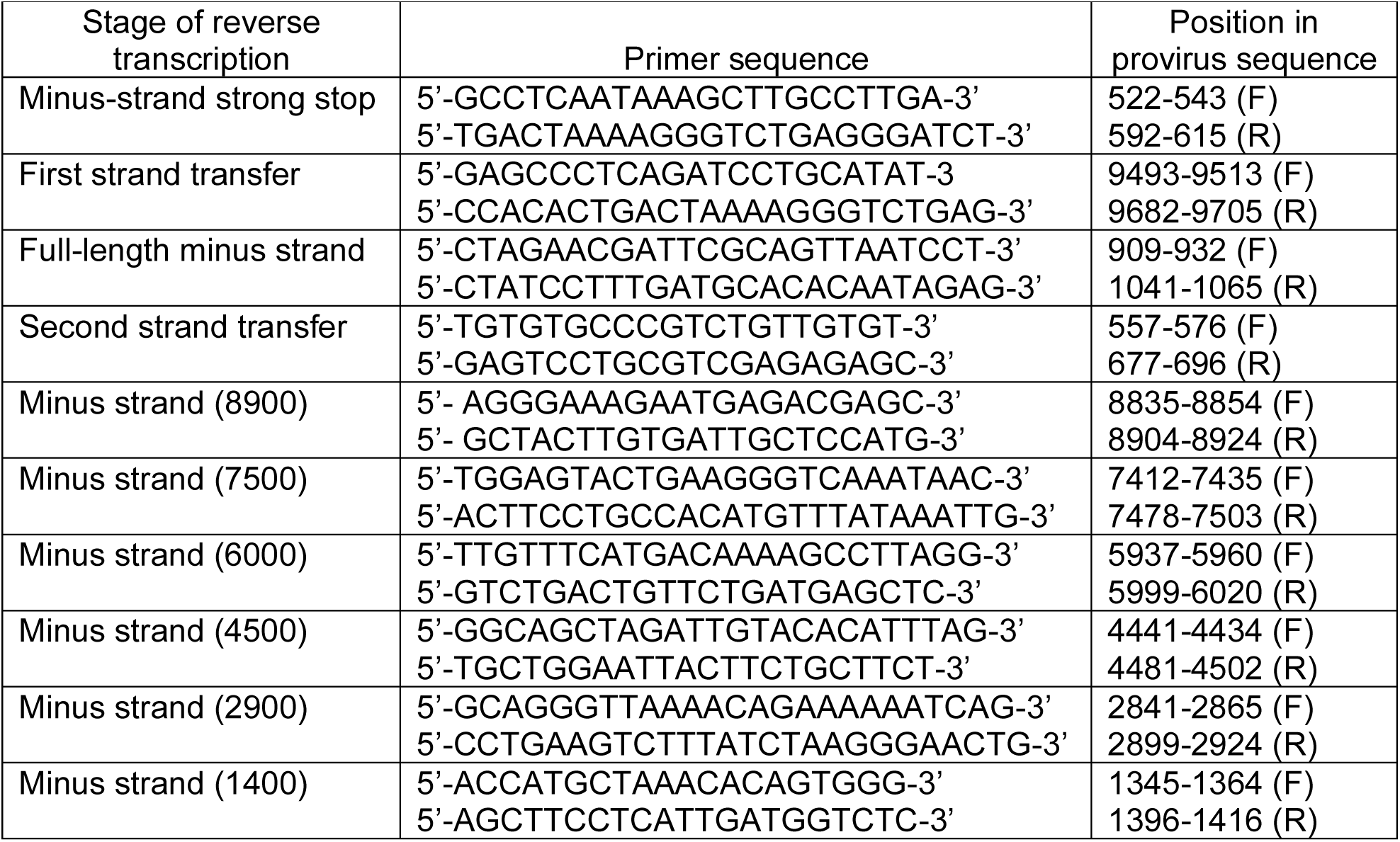
Primers used for quantification of HIV-1 reverse transcripts

Quantitative PCR reactions were performed in 20 μl volumes with a Stratagene Mx3000p real time thermal cycler according to the following program: 95° for 10 min (1 cycle) followed by 95°C for 30s, 55° for 1 min, and 72° for one min (40 cycles). DNA copy numbers were interpolated from standard curves of C_t_ values generated from reactions containing dilutions of R9 proviral plasmid DNA, performed in parallel.

### Uncoating Assays

ERT reactions (50 μl) were incubated at 37°C for 6h, diluted with 450 μl cold reaction buffer lacking dNTPs, and subjected to centrifugation at 100,000 × g in a Beckman TLA-55 rotor. Control reactions were diluted and pelleted immediately after mixing. The supernatants were withdrawn and transferred to clean tubes, and the pellets were resuspended in ERT reaction buffer (0.5 ml). The supernatants and pellets were assayed for CA protein by p24 ELISA (41) and for RT activity using an exogenous primer-template assay, as previously described (42). The resulting values were used to calculate the fraction of pelleted CA and RT activity in the samples.

## Acknowledgments

This work was supported by NIH grants R56 AI076121 and P50 AI150481 (Pittsburgh Center for HIV-Protein Interactions). We thank the staff of the Vanderbilt Cryo-Electron Microscopy Facility for training and facilities for electron microscopy and Dr. Greg Sowd for critical reading of the manuscript.

## Notes

### Competing Interest Statement

The authors have declared no competing interest.

## References

1. Forshey BM, von Schwedler U, Sundquist WI, Aiken C. 2002. Formation of a human immunodeficiency virus type 1 core of optimal stability is crucial for viral replication. J Virol 76:5667–5677.

2. Yufenyuy EL, Aiken C. 2013. The NTD-CTD intersubunit interface plays a critical role in assembly and stabilization of the HIV-1 capsid. Retrovirology 10:29.

3. Dismuke DJ, Aiken C. 2006. Evidence for a functional link between uncoating of the human immunodeficiency virus type 1 core and nuclear import of the viral preintegration complex. J Virol 80:3712–3720.

4. Yang R, Shi J, Byeon IJ, Ahn J, Sheehan JH, Meiler J, Gronenborn AM, Aiken C. 2012. Second-site suppressors of HIV-1 capsid mutations: restoration of intracellular activities without correction of intrinsic capsid stability defects. Retrovirology 9:30.

5. Stremlau M, Perron M, Lee M, Li Y, Song B, Javanbakht H, Diaz-Griffero F, Anderson DJ, Sundquist WI, Sodroski J. 2006. Specific recognition and accelerated uncoating of retroviral capsids by the TRIM5alpha restriction factor. Proc Natl Acad Sci U S A 103:5514–5519.

6. Roa A, Hayashi F, Yang Y, Lienlaf M, Zhou J, Shi J, Watanabe S, Kigawa T, Yokoyama S, Aiken C, Diaz-Griffero F. 2012. RING domain mutations uncouple TRIM5alpha restriction of HIV-1 from inhibition of reverse transcription and acceleration of uncoating. J Virol 86:1717–1727.

7. Hu WS, Hughes SH. 2012. HIV-1 reverse transcription. Cold Spring Harb Perspect Med 2.

8. Dharan A, Bachmann N, Talley S, Zwikelmaier V, Campbell EM. 2020. Nuclear pore blockade reveals that HIV-1 completes reverse transcription and uncoating in the nucleus. Nat Microbiol doi:10.1038/s41564-020-0735-8.

9. Farnet CM, Haseltine WA. 1990. Integration of human immunodeficiency virus type 1 DNA in vitro. Proc Natl Acad Sci U S A 87:4164–4168.

10. Monserrate JP, York JD. 2010. Inositol phosphate synthesis and the nuclear processes they affect. Curr Opin Cell Biol 22:365–373.

11. Shen X, Xiao H, Ranallo R, Wu WH, Wu C. 2003. Modulation of ATP-dependent chromatin-remodeling complexes by inositol polyphosphates. Science 299:112–114.

12. Montpetit B, Thomsen ND, Helmke KJ, Seeliger MA, Berger JM, Weis K. 2011. A conserved mechanism of DEAD-box ATPase activation by nucleoporins and InsP6 in mRNA export. Nature 472:238–242.

13. Alcazar-Roman AR, Tran EJ, Guo S, Wente SR. 2006. Inositol hexakisphosphate and Gle1 activate the DEAD-box protein Dbp5 for nuclear mRNA export. Nat Cell Biol 8:711–716.

14. Brehm MA, Klemm U, Rehbach C, Erdmann N, Kolsek K, Lin H, Aponte-Santamaria C, Grater F, Rauch BH, Riley AM, Mayr GW, Potter BVL, Windhorst S. 2019. Inositol hexakisphosphate increases the size of platelet aggregates. Biochem Pharmacol 161:14–25.

15. Wickner RB, Kelly AC, Bezsonov EE, Edskes HK. 2017. [PSI+] prion propagation is controlled by inositol polyphosphates. Proc Natl Acad Sci U S A 114:E8402–E8410.

16. Wei H, Landgraf D, Wang G, McCarthy MJ. 2018. Inositol polyphosphates contribute to cellular circadian rhythms: Implications for understanding lithium’s molecular mechanism. Cell Signal 44:82–91.

17. Dick RA, Zadrozny KK, Xu C, Schur FKM, Lyddon TD, Ricana CL, Wagner JM, Perilla JR, Ganser-Pornillos BK, Johnson MC, Pornillos O, Vogt VM. 2018. Inositol phosphates are assembly co-factors for HIV-1. Nature 560:509–512.

18. Mallery DL, Marquez CL, McEwan WA, Dickson CF, Jacques DA, Anandapadamanaban M, Bichel K, Towers GJ, Saiardi A, Bocking T, James LC. 2018. IP6 is an HIV pocket factor that prevents capsid collapse and promotes DNA synthesis. Elife 7.

19. Yong WH, Wyman S, Levy JA. 1990. Optimal conditions for synthesizing complementary DNA in the HIV-1 endogenous reverse transcriptase reaction. Aids 4:199–206.

20. Warrilow D, Meredith L, Davis A, Burrell C, Li P, Harrich D. 2008. Cell factors stimulate human immunodeficiency virus type 1 reverse transcription in vitro. J Virol 82:1425–1437.

21. Borroto-Esoda K, Boone LR. 1991. Equine infectious anemia virus and human immunodeficiency virus synthesis in vitro: characterization of the endogenous reverse transcriptase reaction. J Virol 65:1952–1959.

22. Fassati A, Goff SP. 2001. Characterization of intracellular reverse transcription complexes of human immunodeficiency virus type 1. J Virol 75:3626–3635.

23. Chan EW, Dale PJ, Greco IL, Rose JG, O’Connor TE. 1980. Effects of polyethylene glycol on reverse transcriptase and other polymerase activities. Biochim Biophys Acta 606:353–361.

24. Jacques DA, McEwan WA, Hilditch L, Price AJ, Towers GJ, James LC. 2016. HIV-1 uses dynamic capsid pores to import nucleotides and fuel encapsidated DNA synthesis. Nature 536:349–353.

25. Bhattacharya A, Alam SL, Fricke T, Zadrozny K, Sedzicki J, Taylor AB, Demeler B, Pornillos O, Ganser-Pornillos BK, Diaz-Griffero F, Ivanov DN, Yeager M. 2014. Structural basis of HIV-1 capsid recognition by PF74 and CPSF6. Proc Natl Acad Sci U S A 111:18625–18630.

26. Price AJ, Jacques DA, McEwan WA, Fletcher AJ, Essig S, Chin JW, Halambage UD, Aiken C, James LC. 2014. Host cofactors and pharmacologic ligands share an essential interface in HIV-1 capsid that is lost upon disassembly. PLoS Pathog 10:e1004459.

27. Shi J, Zhou J, Shah VB, Aiken C, Whitby K. 2011. Small-molecule inhibition of human immunodeficiency virus type 1 infection by virus capsid destabilization. J Virol 85:542–549.

28. Saito A, Ferhadian D, Sowd GA, Serrao E, Shi J, Halambage UD, Teng S, Soto J, Siddiqui MA, Engelman AN, Aiken C, Yamashita M. 2016. Roles of Capsid-Interacting Host Factors in Multimodal Inhibition of HIV-1 by PF74. J Virol 90:5808–5823.

29. Peng K, Muranyi W, Glass B, Laketa V, Yant SR, Tsai L, Cihlar T, Muller B, Krausslich HG. 2014. Quantitative microscopy of functional HIV post-entry complexes reveals association of replication with the viral capsid. Elife 3:e04114.

30. Balasubramaniam M, Zhou J, Addai A, Martinez P, Pandhare J, Aiken C, Dash C. 2019. PF74 Inhibits HIV-1 Integration by Altering the Composition of the Preintegration Complex. J Virol 93.

31. Bunce CM, French PJ, Allen P, Mountford JC, Moor B, Greaves MF, Michell RH, Brown G. 1993. Comparison of the levels of inositol metabolites in transformed haemopoietic cells and their normal counterparts. Biochem J 289 (Pt 3):667–673.

32. Mallery DL, Faysal KMR, Kleinpeter A, Wilson MSC, Vaysburd M, Fletcher AJ, Novikova M, Bocking T, Freed EO, Saiardi A, James LC. 2019. Cellular IP6 Levels Limit HIV Production while Viruses that Cannot Efficiently Package IP6 Are Attenuated for Infection and Replication. Cell Rep 29:3983–3996 e3984.

33. Hulme AE, Perez O, Hope TJ. 2011. Complementary assays reveal a relationship between HIV-1 uncoating and reverse transcription. Proc Natl Acad Sci U S A 108:9975–9980.

34. Rankovic S, Ramalho R, Aiken C, Rousso I. 2018. PF74 Reinforces the HIV-1 Capsid To Impair Reverse Transcription-Induced Uncoating. J Virol 92.

35. Rankovic S, Varadarajan J, Ramalho R, Aiken C, Rousso I. 2017. Reverse Transcription Mechanically Initiates HIV-1 Capsid Disassembly. J Virol 91.

36. Karageorgos L, Li P, Burrell CJ. 1995. Stepwise Analysis of Reverse Transcription in a Cell-to-Cell Human-Immunodeficiency-Virus Infection Model - Kinetics and Implications. Journal of General Virology 76:1675–1686.

37. Chesebro B, Wehrly K, Nishio J, Perryman S. 1992. Macrophage-tropic human immunodeficiency virus isolates from different patients exhibit unusual V3 envelope sequence homogeneity in comparison with T-cell-tropic isolates: definition of critical amino acids involved in cell tropism. Journal of virology 66:6547–6554.

38. Shah VB, Aiken C. 2011. In vitro uncoating of HIV-1 cores. J Vis Exp doi:10.3791/3384.

39. Durocher Y, Perret S, Kamen A. 2002. High-level and high-throughput recombinant protein production by transient transfection of suspension-growing human 293-EBNA1 cells. Nucleic Acids Res 30:E9.

40. Liu X, Harada S. 2013. DNA isolation from mammalian samples. Curr Protoc Mol Biol Chapter 2:Unit2 14.

41. Wehrly K, Chesebro B. 1997. p24 antigen capture assay for quantification of human immunodeficiency virus using readily available inexpensive reagents. Methods 12:288–293.

42. Aiken C. 1997. Pseudotyping human immunodeficiency virus type 1 (HIV-1) by the glycoprotein of vesicular stomatitis virus targets HIV-1 entry to an endocytic pathway and suppresses both the requirement for Nef and the sensitivity to cyclosporin A. Journal of Virology 71:5871–5877.

